# Contribution of genetic variation and developmental stage to methylome dynamics in myeloid differentiation

**DOI:** 10.1101/132985

**Authors:** Xiang Chen, Yiping Fan, Jinjun Cheng, Beisi Xu, Yong-Dong Wang, Donald Yergeau, John Easton, James R. Downing, Jinghui Zhang

## Abstract

DNA methylation is important to establish a cell’s developmental identity. It also modulates cellular responses to endogenous developmental stimuli or environmental changes. We designed an *in vitro* myeloid differentiation model to analyze the genetic and developmental contribution to methylome dynamics using whole-genome bisulfide sequencing and transcriptome sequencing. Using a recursive partitioning approach, we identified 34,502 differentially methylated regions (DMRs) associated with genetic background and/or developmental stimuli. Specifically, 23,792 DMRs (69%) were significantly associated with inter-individual variations, of which 82% were associated with genetic polymorphisms in *cis*. Notably, inter-individual variations further modified 57 of 212 (26%) developmental DMRs with transcriptomic responses. Our study presents a novel analytical approach to determine the bona fide genetic contribution embedded in outlier patterns of CpG-SNPs in individual methylomes. This approach can be used to study genetic and epigenetic mechanisms underlying differential responses to developmental stimuli, environmental changes, and inter-individual differences in drug responses.

## Introduction

Differentiation and development from a single zygote to a multicellular complex organism requires the precise regulation of gene expression at various developmental stages. The complex regulatory system that drives developmental progression depends on the interaction of multiple components, including transcription factor expression and its DNA occupancy(Spitz and Furlong 2012), epigenetic regulation(Reik 2007), and mechanisms of post-translational modification of proteins(Deribe et al. 2010). In addition to modulating developmental changes, epigenetic regulation, including DNA methylation, contributes to phenotype variability among normal individuals. Although differential methylation patterns have been well studied in normal development(Smith and Meissner 2013), diseases(Lopez-Serra and Esteller 2012; Stricker et al. 2013; Schoofs et al. 2014), in different tissues(Ziller et al. 2013), and human populations(Heyn et al. 2013), the effects of inter-individual variability(Jiang et al. 2015) in responses of the methylome (i.e., the set of nucleic acid methylation modifications) to the same developmental stimuli remain largely unexplored.

The role of genetic variations in modulating the DNA methylation pattern has been recognized since the discovery of single-nucleotide polymorphisms (SNPs) at cytosine–phosphate–guanine dinucleotides (CpG-SNPs)(Moser et al. 2009; Shoemaker et al. 2010). Whereas most CpG sites are methylated in mammalian genomes, CpH (H represents A, C, or T) sites are unmethylated(Ehrlich et al. 1982). Therefore, genetic variations at CpG-SNPs are expected to have one allele from a CpG site and the other from a CpH site(Shoemaker et al. 2010). Individuals with CpG/CpG, CpG/CpH, or CpH/CpH genotypes at CpG-SNPs exhibit full methylation, partial methylation, and no methylation, respectively. Therefore, differential methylation at CpG-SNPs is generally considered to be a technical artifact in methylome analysis, and exclusion of CpGs located within 10 bp of known polymorphisms in the dbSNP database is recommended for methylation arrays(Price et al. 2013). However, this filtering strategy also leads to considerable loss (up to one tenth) of CpGs that might carry bona fide variations. Genomewide DNA methylation quantitative trait loci (mQTLs) analyses have established that the DNA methylation could be affected by adjacent genetic variants(Shoemaker et al. 2010; Zhang et al. 2010; Heyn et al. 2014; Zhang et al. 2014). While these studies revealed a fundamental connection between sequence variants and epigenetic differences (differential DNA methylation), they largely relied on array-based methylome measurement platforms and SNP arrays, which have focused on association of differential methylation of single CpG probes on the methylation array with common SNPs. However, methylome is jointly established and maintained by the DNA methyltransferases and the tet methylcytosine dioxygenases, which have local coordinated activities and lead to similar methylation status among adjacent CpGs on the same DNA molecule(Guo et al. 2017). Functionally, gene silencing through DNA methylation involves the hypermethylation of entire CpG islands instead of individual important CpGs(Esteller 2008). Gene expression status is determined by the regional methylation density of a *cis*-element and not that of individual critical CpG sites(Weber et al. 2007). Subsequently, a block structure was proposed for allele-specific methylation patterns(Shoemaker et al. 2010). We hypothesized that a subset of genetic variations can affect methylation levels of multiple CpGs in their adjacent regions, which we termed as regional methylation quantitative trait loci in *cis* (*cisR-*mQTLs). On the other side, although array studies show that 0.16–1.5% of common SNPs can affect adjacent methylation patterns based on single CpG probe analyses(Kerkel et al. 2008; Schalkwyk et al. 2010), the effects of *cisR-*mQTLs associated with private and common SNPs have not been quantified(Schultz et al. 2015), in part because of the technical limitations of array-based platforms.

DNA methylation profiling has transitioned from single-gene–based(Keshet et al. 1985) and array-based (Price et al. 2013) analyses to genome-wide investigation using whole-genome bisulfite sequencing (WGBS)(Lister et al. 2009). WGBS has 2 inherent advantages. First, it measures genome-wide methylation in an unbiased manner, including intergenic and intronic methylations that might regulate transcription and splicing activities. Second, genome-wide genetic variations in a sample, including private SNPs, can be directly inferred from WGBS, which enables investigations of individual-specific SNPs and *cisR*-mQTLs in addition to those correlated with common SNPs.

In this study, we used WGBS and mRNA-sequencing (mRNA-Seq) to analyze unbiased genome-wide DNA methylation and transcriptome patterns of myeloid populations from 3 individuals collected at 3 developmental stages: bone marrow CD34^+^ progenitor cells, early immature myeloid lineage cells, and late mature myeloid cells. By using this *in vitro* myeloid differentiation model, we investigated (1) the relative contribution of inter-individual variations and developmental stimuli on global DNA methylation patterns in the form of differentially methylation regions (DMRs) and (2) the extent of *cisR-*mQTL in healthy human subjects and (3) how various types of DMRs affect gene transcription.

## Results

### In vitro myeloid differentiation model

The highly reproducible hematopoietic cell lineages culture and *in vitro* differentiation model(Gupta et al. 2014) was chosen for our study because it allowed the purification of cells at distinct developmental stages by flow cytometry.

Expression profiles of cell surface markers from *in vitro* myeloid differentiation cultures were analyzed by flow cytometry. CD34^+^ cells in culture gradually differentiated into myeloid lineage cells, as seen by increasing CD11b and CD13 expressions and decreasing CD34 expression over the 12-day period (Figure 1A). The morphology and immunophenotype of cell populations were monitored at day 0, 18 h, day 2, day 3, day 4, day 6, day 9, and day 12. The day 0 population represented bone marrow CD34^+^ progenitor cells that did not express specific lineage surface antigens, the day 3 population represented early immature myeloid lineage cells that had a gradual increase in surface CD13 expression and a decrease in CD34 expression, and the day 12 population represented mature myeloid cells as they had more than 70% neutrophilic granulocytes (Figure 1A and 1B)(Gupta et al. 2014).

**Figure 1.**
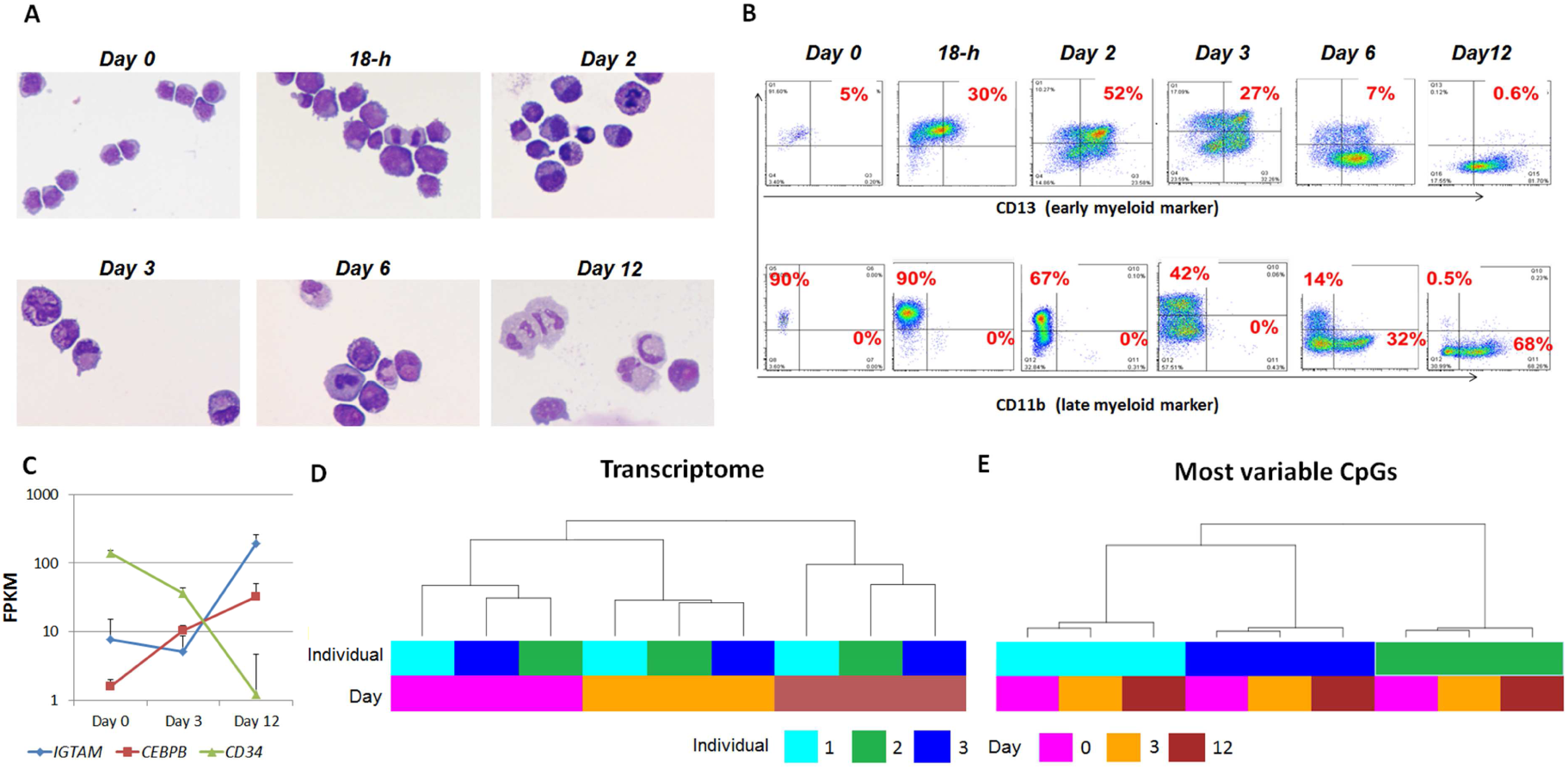
*In vitro* myeloid differentiation experiment. (A) Wright–Giemsa staining of cytospin preparations at different time points during in vitro myeloid differentiation. (B) Immunophenotype analysis by flow cytometry at different time points during in vitro myeloid differentiation. (C) Expression of representative genes *ITGAM*, *CEBPE*, and *CD34*. (Error bar represents 1 standard deviation across the 3 individuals.) (D) Hierarchical clustering analysis of transcriptome data. (E) Hierarchical clustering analysis of the most variable CpGs. FPKM, fragments per kilo base per million.

### Genome-wide transcriptome and methylome profiles in bone marrow–derived myeloid cells

WGBS and mRNA-Seq were performed, respectively, on DNA and RNA extracted from 9 samples obtained from *in vitro*–differentiated bone marrow cells. On average, WGBS sequenced 133.1 billion nucleotides (range 123.2–141.3) per sample, with 111.7 billion nucleotides (range 99.9–120.7) being uniquely mapped (average depth 33×). The average genome-wide bisulfite conversion rate was 0.995 (range 0.993–0.997), and 80.0% (range 75.4%–85.8%) of CpGs were covered at ≥5× (Supplementary Table 1)(Ziller et al. 2015). Further, 9.77 billion nucleotides (range 9.16–10.34) were sequenced per RNA-Seq sample, of which 8.86 billion (range 8.28– 9.23) were uniquely mapped. On average, 49.1% of exonic bases were covered at ≥20× (range 47.3–50.2, Supplementary Table 2).

Expressions profiles of selected neutrophil-specific genes such as *CD34*, *ITGAM* and the myeloid transcriptional regulator *CEBPE* were studied to verify the *in vitro* myeloid differentiation model. Notably, a decrease in *CD34* expression was accompanied by an increase in *CEBPE* expression throughout differentiation (Figure 1C). In contrast, *ITGAM* expression was largely unchanged from progenitor cells to early myeloid cells, but increased dramatically in late myeloid cells. Therefore, the expression profiles of selected genes confirmed the gradual increase in the extent of myeloid maturation(Nakajima and Ihle 2001).

Overall, 10,308 of 27,807 annotated genes were differentially expressed genes (DEGs), and they clustered according to the stage of myeloid developmental (Figure 1D). Consistent with this profile, 9120 DEGs (88.5%) showed a significant developmental difference (false discovery rate [FDR] *q-*value ≤ 0.05) (Yuan et al. 2007). In contrast, only 74 (0.1%) transcripts showed significant differences among the 3 individuals (i.e., inter-individual variations), suggesting a minimal contribution from inter-individual variations at the transcriptome level. The remaining DEGs (11.4%) were largely stochastic, without showing a significant developmental or inter-individual difference. Interestingly, non-coding transcripts (*n*=21) were significantly enriched in genes that had inter-individual variations (*P*=0.013, odds ratio=1.94, Fisher exact test). This result was consistent with that of a recent report showing higher inter-individual expression variability in non-coding RNAs than in protein-coding RNAs(Kornienko et al. 2016). For example, the expression of lincRNA AC104135.3 was stable across developmental stages but had high inter-individual variation (average fragments per kilo base per million [FPKM] in 3 individuals: 1.51±0.03, 3.11±0.52, and 0.009±0.004, respectively; *q*-value = 0.0038). The miRNA expression followed a similar pattern, because differential expression was largely driven by transitions through developmental stages (Supplementary Figure 1).

### Genetic variations contribute to inter-individual variations in methylation profiles

The 14.3 million CpGs covered at ≥5× in the WGBS analysis(Ziller et al. 2015) were used to analyze inter-individual and developmental variations in the methylation profile at each CpG site. The global methylation profile showed a bimodal distribution, with very similar distribution patterns among the 3 individuals (Supplementary Figure 2A). There was a trend toward hypomethylation in the global profile during the myeloid differentiation process, especially between the early myeloid stage (day 3) and the late myeloid stage (day 12) (Supplementary Figure 2B).

**Figure 2.**
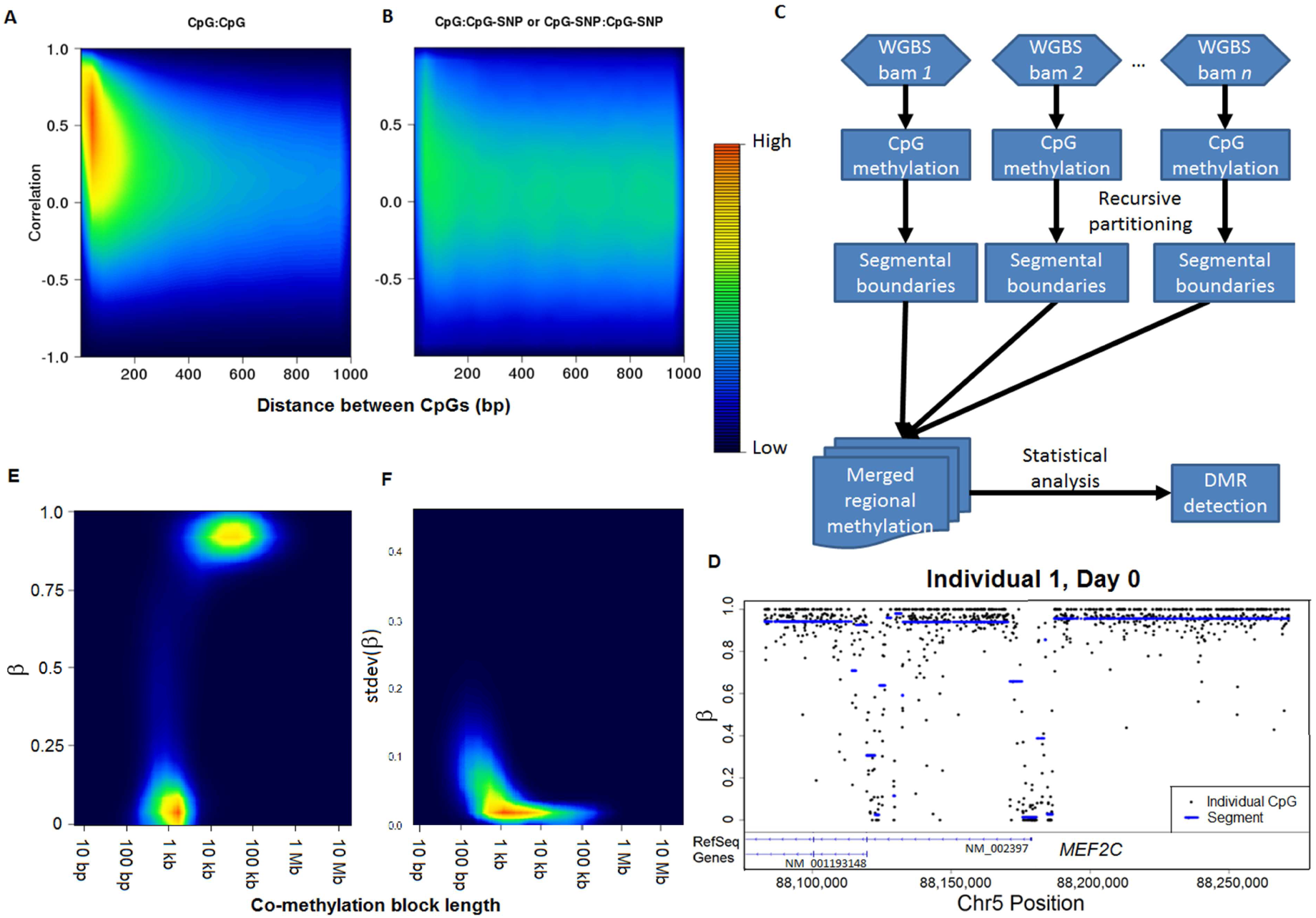
Co-methylation blocks and outlier methylation patterns in CpG-SNPs. (A) Positive correlation of methylation status in neighboring CpGs. Density was calculated as the frequency of CpG pairs that fall in an area in the scatter plot (brighter red at a higher frequency). (B) Absence of overall correlation of methylation status between CpG-SNPs and adjacent CpGs. (C) Pipeline for regional methylation analysis. (D) Example of recursive partitioning–based segmentation of WGBS data. (E) Single-sample segmentation showing a capture of spatial distribution of the human methylome. (F) Enrichment of methylation variations in small blocks. Regions were defined by pooling segmental boundaries across all samples (see subpart C), and the cross-sample variation of segmental methylation levels was calculated. stdev, standard deviation; WGBS, whole-genome bisulfite sequencing; DMR, differentially methylated region.

Overall, only 181,409 (1.27%) of CpG sites showed significant differential methylation. In contrast to gene expression, hierarchical clustering using the 50,000 most variable CpGs was grouped by individuals instead of developmental stage (Figure 1E). Specifically, 180,419 CpGs (99.5% of all CpGs with differential methylation) showed significant inter-individual differences in methylation (*q* value ≤ 0.05), compared to only 993 CpGs showing significant development-related differences. To determine the contribution from common SNPs, 2 filtering strategies were used: (1) CpGs overlapping with dbSNPs were filtered (2.2% of all CpGs), or (2) CpGs located within 10 bp of common dbSNPs were filtered (additional 10.7% of all CpGs). Inter-individual differences in clustering patterns persisted in both strategies (Supplementary Data and Supplementary Figure 3), suggesting that differential methylation can be affected by private genetic variations that are not found in dbSNPs. Indeed, an additional 262,367 private CpG-SNPs were identified by WGBS analysis. Altogether, CpG-SNPs, which include both dbSNPs and private SNPs, strongly correlated with the CpG allele count (average coefficient 0.425, range 0.413–0.439, *P*<2.2×10^−16^, Supplementary Figure 4) and accounted for 68.7% of CpGs exhibiting significant inter-individual differences. The removal of CpG-SNPs (Supplementary Figure 5A) led to a cluster of samples at the late myeloid stage. Inter-individual variations remained predominant in the progenitor and early myeloid stages.

### Recursive partitioning reveals block structure of methylome architecture

Our results showed that differentially methylated CpG sites located within 150 bp were highly correlated (Figure 2A), supporting the presence of a block structure that is consistent with a previous report(Shoemaker et al. 2010). However, this structure was not seen when only CpG-SNPs were used (Figure 2B), suggesting that CpG-SNPs themselves are outliers that cannot be used to assess biologically driven methylation changes (Supplementary Figure 6).

To support the use of methylation blocks for robust analysis, an analytical process that used a recursive segmentation approach to define block structures that exhibit co-methylation patterns was developed (Figure 2C). Specifically, each sample was first partitioned by using the regression-tree approach into a set of blocks on the basis of methylation transition patterns (Figure 2D). Breakpoints across all samples were then pooled to derive a unified set of regions to reveal differentially methylated regions (DMRs). This analytical approach focuses on co-methylation block structures and identifies blocks with differential methylation levels among samples, whereas outlier methylation patterns from CpG-SNPs are diluted in blocks and less likely to affect the analysis. This approach was used to study the contribution of inter-individual variations and developmental stimuli to methylation changes observed in the in vitro myeloid differentiation model.

Results from single-sample segmentation showed that the segmentation approach faithfully captured spatial distribution of the human methylome, in which most CpG sites are heavily methylated whereas CpG islands remain unmethylated (Figure 2E). There were 2 major clusters of blocks: (1) a set of unmethylated blocks approximately 1 kb in length, which is consistent with the average length of CpG islands(Deaton and Bird 2011), and (2) a set of fully methylated blocks of a larger size (10–100 kb). For example, CpGs in regions surrounding the canonical transcription start site (TSS) of *MEF2C* (NM_002397) as well as an alternative TSS (NM_001193148) were hypomethylated, whereas intergeneic and most of the intronic regions were fully methylated with stochastic fluctuations. The segmentation approach accurately captured this pattern and was largely robust to intrinsic noises (Figure 2D). An evaluation of methylation variations across all samples revealed that large blocks (>10 kb) were absent when only blocks that showed differential methylation during *in vitro* myeloid differentiation were considered (Figure 2F), suggesting that changes in the methylome were focal during *in vitro* myeloid differentiation. Clustering of blocks with the most variable methylation regions (Supplementary Figure 5B) revealed a signature nearly identical to that seen in the CpG-SNP filtered single CpG analysis (Supplementary Figure 5A). These consistent results confirm that the recursive partitioning–based co-methylation analysis is robust to outlier effects from CpG-SNPs and accurately captures the embedded differential methylation patterns.

Three types of DMRs were identified: DMRs associated with variations among normal individuals (i-DMRs), DMRs associated with developmental stimuli (d-DMRs), and DMRs for which methylation responses to developmental stimuli are dependent on inter-individual variations (di-DMRs). An example of i-DMR was a 4-kb region spanning the exon–intron junctions with >100 CpG sites in *MUM1*. The region was hypomethylated in individual 3 compared with individuals 1 and 2, which correlates with the genotype of rs80117987 (chr19:1367095, GG genotype in individuals 1 and 2 and AG in individual 3). Methylation levels in all 3 individuals remained unchanged through the 3 developmental stages (Figure 3A). An example of d-DMR was a 2.8-kb region in *SYMPK*, which was hypermethylated in all 3 individuals in the mature myeloid stage (day 12) but not the progenitor (day 0) or the early myeloid cell (day 3) stages (Figure 3B). A 2.4-kb intronic region on *FOXK1* showed a methylation pattern defined by a di-DMR. In this case, hypermethylation occurred in individual 1 at the progenitor stage (day 0) but not at the early myeloid (day 3) and mature myeloid (day 12) stages. Methylation was not observed in individuals 2 and 3 at any developmental stage (Figure 3C). Overall, 34,502 DMRs were identified: 17,483 i-DMRs, 10,710 d-DMRs, and 6309 di-DMRs (Figure 3D). Intergentic regions were significantly enriched in i-DMRs compared to d-DMRs and di-DMRs (*p* < 2.2 x 10^−16^, OR = 1.36 and *p* = 2.3 x 10^−13^, OR = 1.37 by Fisher’s exact test, respectively).

**Figure 3.**
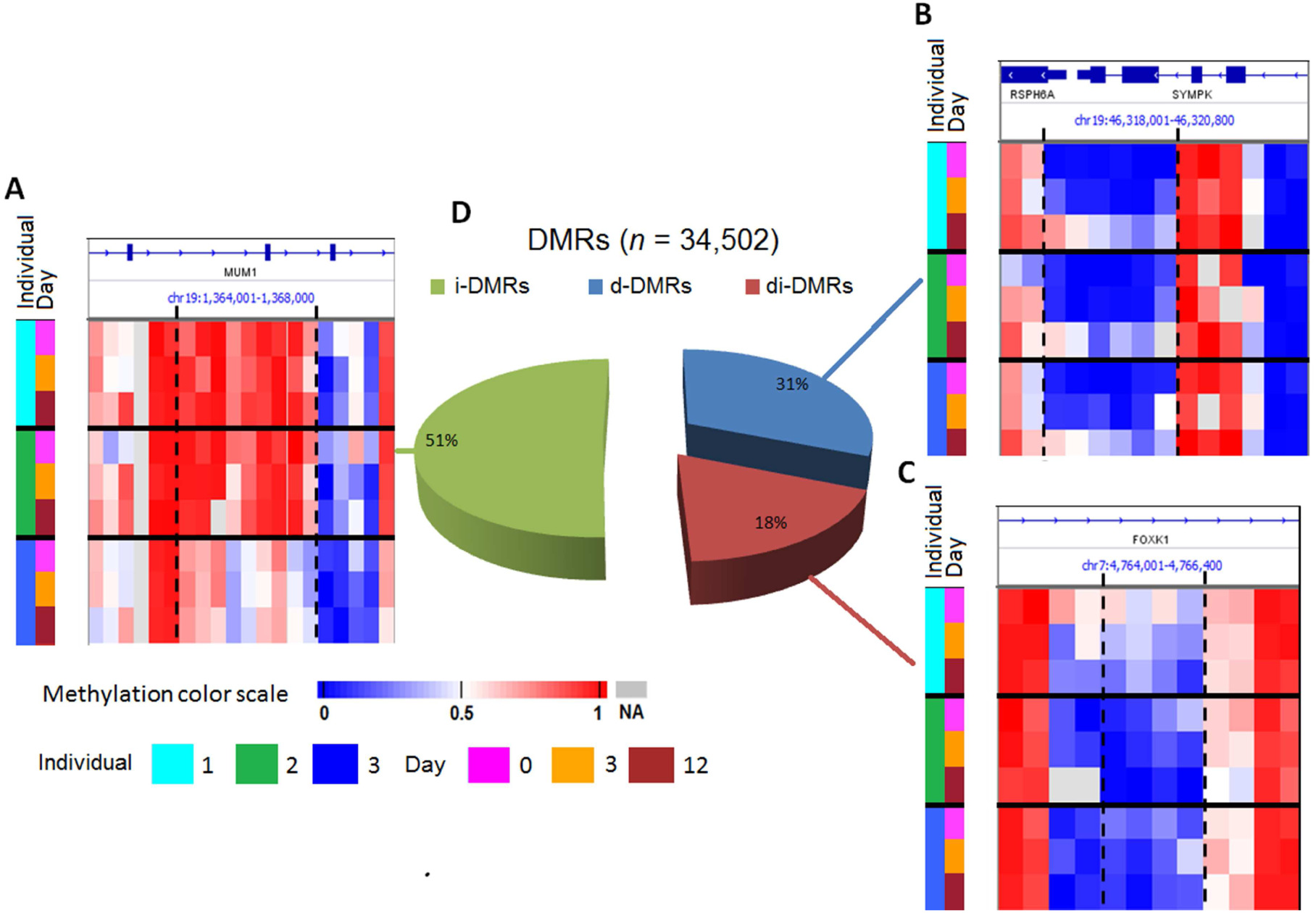
Differentially methylated regions identified in the in vitro myeloid differentiation model. (A) An i-DMR in *MUM1*. (B) A d-DMR in *SYMPK*. (C) An id-DMR in *FOXK1*. (D) Pie-chart showing the types of DMRs. DMR, differentially methylated region; NA: data not available due to lack of sufficient coverage

A joint analysis of the methylation level of i-DMRs and di-DMRs with adjacent SNPs showed 82% of the i-DMRs and di-DMRs were associated with the genotype of adjacent SNPs, which forms the *cisR-*mQTL pattern.

### Inter-individual variations affect developmental responses

Because DNA methylation is a common mammalian epigenetic mechanism regulating chromatin accessibility and gene transcription,(Bird 2002) we investigated the correlation pattern between gene expression levels and regional methylation levels (from 10 kb upstream of the TSS to the transcription end site [TES]). A total of 227 DMRs were significantly correlated with the expression of 212 *cis*-genes. Tracking with the transcriptomic pattern, 96% of DMRs regulating *cis-*gene expression were d-DMRs (*n*=162) or di-DMRs (*n*=57).

Figure 4 shows an example of a di-DMR (chr4:185734543-185735006) in the first intron of *ACSL1*. This region harbors many potential active transcription factor binding sites in the human myeloid leukemia cell line K562 (Figure 4A). The di-DMR was hypomethylated during differentiation, which correlates with the upregulation of *ACSL1* expression (Figure 4B). This region also exhibited differential methylation levels, with a higher methylation seen in individual 1 on days 3 and days 12 than for individuals 2 and 3. Interestingly, at rs116679280, individual 1 had a heterozygous A/G genotype whereas individuals 2 and 3 had a homozygous G/G genotype. Consequently, although the baseline expression level of *ACSL1* in progenitor cells (day 0) was similar among all 3 individuals, its expression level at the mature myeloid stage (day 12) in individual 1 was 36%–42% lower than that in individuals 2 and 3 (Figure 4B). The allelic expression pattern of *ACSL1* in each individual was also evaluated, using heterozygous variations inferred from WGBS data. Specifically, the allelic-specific expression (ASE) score was calculated for the samples. The ASE score was high in only individual 1 (Figure 4C), further suggesting that the inter-individual differences in methylation caused by genetic variations may result in differential gene expression in response to developmental stimuli. Overall, inter-individual variations modified methylation signatures in 26% of the functional DMRs regulated by differentiation stimuli in our *in vitro* myeloid differentiation model.

**Figure 4.**
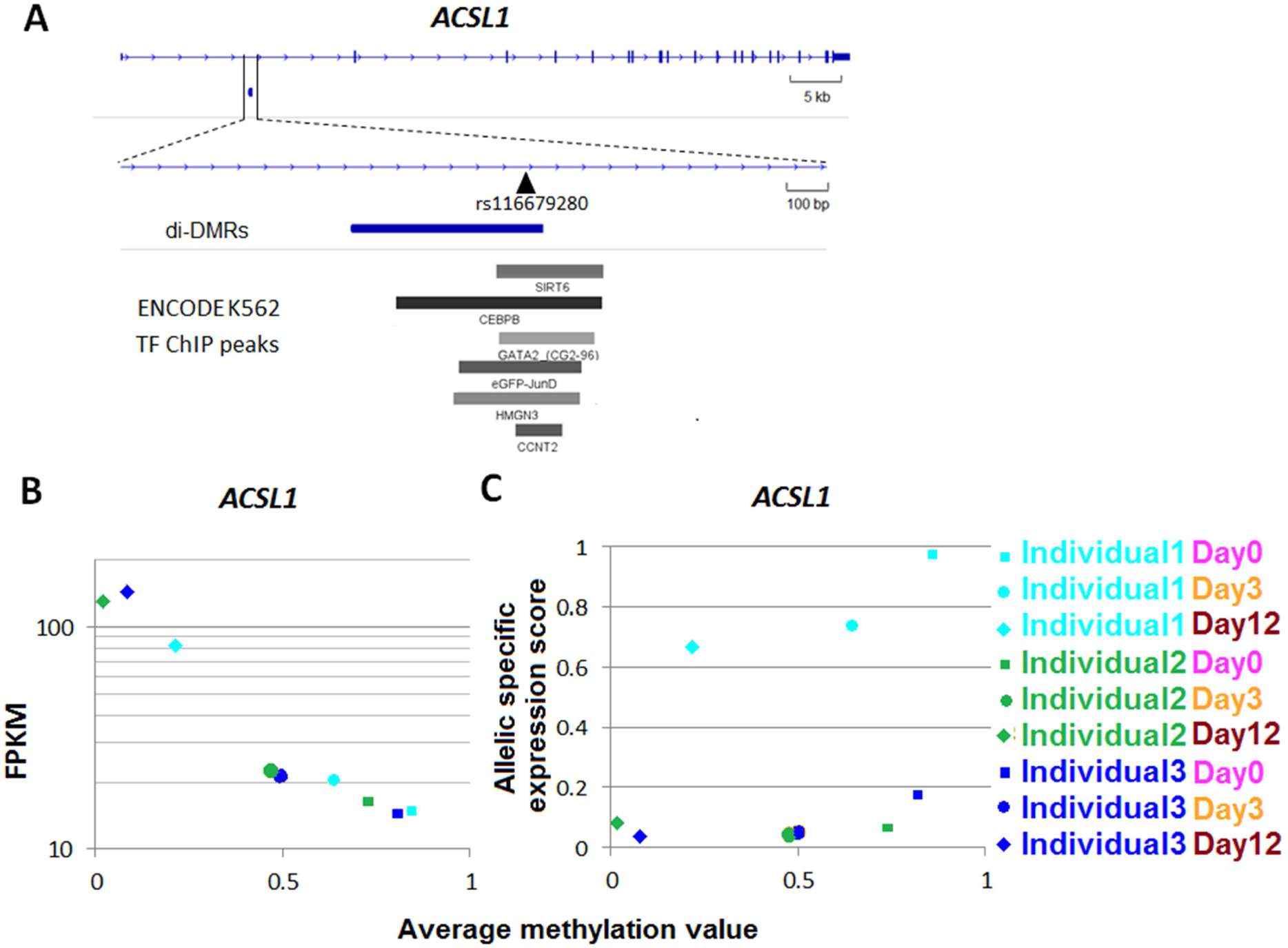
di-DMRs correlate with differential gene expression.(A) Structure of the *ACSL1* gene and di-DMR (a *cisR-*mQTL associated with rs116679280). (B) *ACSL1* expression is correlated with the methylation level of id-DMR. (C) ASE score of *ACSL1* was consistently identified in individual 1. FPKM, fragments per kilo base per million; DMR, differentially methylated region; ASE, allele-specific expression.

## Discussion

The epigenome, which describes the set of complex chemical modifications associated with DNA and histones, instructs gene expression and determines the developmental identity of each cell. It is characterized by a dynamic response to developmental stimuli such as cytokines(Deverman and Patterson 2009), growth factors,(Lovicu et al. 2011) and hormones(Dohler and Wuttke 1976). The genomic and epigenomic signatures of individuals jointly determine their responses to developmental changes.

Although epigenetic responses to various developmental and environmental changes have been systematically studied throughout the human lifespan(Kanherkar et al. 2014) and in various disease states(Lopez-Serra and Esteller 2012; Stricker et al. 2013; Schoofs et al. 2014), the extent to which genetic profiles contribute to shaping an individual’s epigenome remains unclear. To investigate this important yet understudied mechanism, we designed an *in vitro* myeloid differentiation model and a recursive partitioning–based segmentation approach to investigate the joint contribution of developmental stage and genetic variations to the methylation dynamics. Our results show that genetic background is a major contributor to shaping an individual’s methylome architecture for a single cell in differentiation. In our model, 51% of all DMRs were attributed to inter-individual variations alone (i-DMRs), and an additional 18% of DMRs were attributed to joint effects from inter-individual variations and developmental stimuli (di-DMRs). Consistent with previous reports based on single CpG probes(Zhi et al. 2013), our analysis also revealed that 82% of i-DMRs and di-DMRs formed *cisR-*mQTLs, and the remaining DMRs might be associated with other genetic variations (e.g., small insertions and deletions or microsatellites) not evaluated in this study. Nevertheless, the high percentage of *cisR-*mQLTS in i-DMRs and di-DMRs emphasizes the important role of genetic background in defining an individual’s methylome. The abundance of di-DMRs further indicates that the genetic background of individuals plays important roles in fine tuning their developmental responses at the level of DNA methylation. Moreover, 26% of developmentally regulated DMRs with roles in *cis-*gene regulation were di-DMRs, suggesting that DNA methylation is the major mechanism by which the genetic background modulates an individual’s transcriptomic response to external stimuli.

WGBS methylome analysis can be generally grouped into single CpG-based analyses and co-methylation–based regional analyses. Compared with other methylation platforms, WGBS is relatively inefficient because approximately 80% of the total reads are non-informative for analyzing CpG methylation(Ziller et al. 2013). Thus, a recent analysis suggested that methylomes generated at a 30× coverage are not adequate for single CpG-based differential methylation analysis, but co-methylation analysis based on regional approaches can partially recover the lost information(Libertini et al. 2016). The recursive partitioning technique is an efficient nonparametric statistical learning method with a wide range of applications in bioinformatics, such as classification of gene expression, analysis of protein–protein interactions, discovery of biomarkers, and statistical genomics(Chen et al. 2011; Qi 2012). The technique is also powerful in identifying genome-wide regional segmentation patterns(Olshen et al. 2004; Chen et al. 2015). Here, we adapted and improved the recursive partitioning algorithm to analyze regional co-methylation patterns. The approach uses the segmentation algorithm^13^ to derive blocks of co-methylation structures in methylomes and then combine these regions across different samples in the dataset. The segmentation-based regional methylation approach is advantageous for methylome analysis because (1) it is adaptive to the segmental length or number of CpGs in the segments (i.e., no fixed window size), and (2) unlike hidden Markov model–based approaches(Stadler et al. 2011), it does not require prior knowledge of the potential number of methylation states. The application of our recursive partitioning–based segmentation approach to analyze data from our *in vitro* myeloid differentiation model suggested that the algorithm identifies the characteristic features of human methylomes (e.g., short blocks of unmethylated CpG islands dispersed among long stretches of fully methylated regions). Although the breakpoints pooling across multiple samples resulted in a reduced block length, a large fraction of long blocks was retained. Our analysis suggests that our model is not only robust to methylation outliers caused by CpG-SNPs but also powerful for detecting DMRs.

Our study design used an *in vitro* model to control development stimuli, which are critical to evaluate analytical approaches for investigating methylation changes driven by genetic and developmental transitions. The sample size of this study was small, and additional DMRs might be discovered in larger cohorts. For example, some large blocks with partial methylation can be broken into smaller regions for larger sample sizes. Nevertheless, our study presents a novel approach to determine the contribution of individual genetic variations during development. From a clinical standpoint, our study offers new insights for analyzing large-scale WGBS data in ongoing cancer studies(Kulis et al. 2015).

## Methods

### Bone marrow–derived myeloid cell cultures

Human bone marrow–derived CD34^+^ progenitor cells were obtained through the Lonza Research Bone Marrow Program (1M-101C, Lonza). Cells were isolated from 3 healthy female African American individuals (age 22–25 years) to reduce potential confounding effects. Bone marrow– derived CD34^+^ cells were isolated by positive immunomagnetic selection from the mononuclear fraction, and a cell purity of ≥90% was confirmed by flow cytometry. Bone marrow CD34^+^ cells for each culture experiment were isolated from a single individual, and the experiment was independently repeated 3 times.

Primary CD34^+^ bone marrow progenitor cells were cultured with a combination of cytokines and monitored for myeloid differentiation *in vitro* to obtain representative and homogeneous populations of early and late myeloid cells. Briefly, freshly harvested cells shipped at an ambient temperature overnight from the vendor were designated as the day 0 sample. Immediately after the cells were received, a small aliquot was used for cytospin slide preparations and Wright– Giemsa staining of cells for flow cytometry analysis. Approximately 0.3 million CD34^+^ cells were collected for RNA or DNA extraction (day 0 sample). All remaining cells were cultured in Iscove’s Modified Dulbecco’s Medium (IMDM) supplemented with 10% fetal calf serum (Thermo Fisher Scientific), 100 ng/mL recombinant human stem cell factor, and 100 ng/mL recombinant human interleukin-3 (PeproTech), with a density of 0.1 million cells per milliliter of medium. On the third day, cells were harvested for phenotype analysis, and 0.3 million cells (day 3 sample) were collected for RNA and DNA extraction. The remaining cells were washed and resuspended in IMDM with 100 ng/mL IL-3, 100 ng/mL stem cell factor, and 10 ng/mL granulocyte colony-stimulating factor (PeproTech). The medium was changed every 2 days, and cells were harvested on day 12 for phenotypic analysis and RNA and DNA extraction (day 12 sample).

Harvested cells were stained with fluorescence-conjugated anti-human CD34, CD13, and CD11b monoclonal antibodies (BD Biosciences Pharmingen, 560941, 557454, and 561685) and analyzed by flow cytometry on a BD LSRII or BD FACSAriaII (BD Biosciences). Data were analyzed using the FlowJo software (Tree Star, Inc.).

### miRNA array, RNA sequencing, and WGBS data generation

In vitro–cultured bone marrow cells at different time points were collected, cryopreserved, and thawed for DNA and RNA isolation by using RNeasy and DNeasy, respectively (QIAGEN). DNA and RNA were quantitated by the Qubit fluorometer and stored at −80°C for further processing.

To construct RNA-Seq libraries, 2–5 μg of total RNA was extracted from samples by using the RNeasy Mini Kit (QIAGEN) according to the manufacturer’s instructions. RNA integrity was measured by using an 2100 Bioanalyzer Lab-on-a-Chip platform (Agilent Technologies). Total RNA was treated with DNase I (Invitrogen) and enriched for poly A–containing mRNA by using Dynabeads Oligo (Invitrogen). cDNA synthesis was performed by using random hexamers and the Superscript Double-Stranded cDNA Synthesis Kit (Invitrogen).

Total RNAs were labeled by using the miRNA Complete Labeling and Hyb Kit (Agilent Technologies), followed by hybridizing to the Human miRNA v19 Microarray (Agilent Technologies, 046064), which contains 4774 unique biologic featured probes targeting 2006 mature miRNAs according to human miRBase, version 19.0 (www.mirbase.org, August 2012).

Microarrays were scanned by using an G2565CA Array Scanner System (Agilent Technologies) at a resolution of 3 μm, and data were extracted by the Feature Extraction software (v10.5.1.1) (Agilent Technologies), using the miRNA_107_Sep09 protocol. Data were processed by the Partek Genomics Suite 6.5 (Partek Inc). After quantile normalization among arrays, each probe was summarized with a single log intensity value.

To construct WGBS libraries, DNAs were sheared by sonication with a E220 Focused-ultrasonicator (Covaris), and DNA fragments were enriched to a fragment size of approximately 300 bp by using AMPure XP beads (Agencourt Bioscience Corp.). After end repair and adenylation, the methylation adaptors (Illumina, Part 100-0010) were ligated by using DNA T4 ligase (NEB, M0202L) for 1 h at 16°C. Bisulfite conversion of genomic DNA was performed by using the EZ DNA Methylation Kit (Zymo Research) according to the manufacturer’s protocol. Briefly, 0.8–1.2 μg of DNA was denatured at 95°C for 30 sec and bisulfite conversion was carried out at 64°C for 2.5 h, followed by another cycle of denaturation at 95°C for 30 sec and bisulfite conversion at 55°C for 15 min. After conversion, samples were desulphonated and eluted by using a column preparation. Adaptor-ligated DNA was enriched through 8 cycles of PCR by using the KAPA HiFi Hotstart Uracil+ kit (KAPA Biosystems). PCR products were purified by AMPure XP beads, and the final fragment size was enriched to 400–450 bp before loading on to a flow cell for clustering and for sequencing (Illumina HiSeq 2500).

### RNA-seq data analysis

Paired-end reads from mRNA-Seq were aligned to the following 4 database files by using a Burrows–Wheeler Aligner (0.5.5): (i) human NCBI Build 37 reference sequence, (ii) RefSeq, (iii) a sequence file representing all possible combinations of non-sequential pairs in RefSeq exons, and (iv) AceView flat file downloaded from UCSC, representing transcripts constructed from a human expressed sequence tag. The final BAM (compressed binary version of the Sequence Alignment/Map [SAM] format) file was constructed by selecting the best alignment in the 4 databases.

Contributions of inter-individual differences and developmental differentiation to gene expression variations (FPKM ≥ 1 in the sample with highest expression, ≥2-fold difference between the samples with the highest and lowest expressions) were evaluated in a linear regression model (categorical variables for both subject identifiers (IDs) and developmental stages):

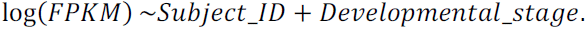

Stepwise backward model selection was performed by using the *step()* function in R (version 3.0.1).

Hierarchical clustering was performed by using the Spearman rank correlation distance for RNA-Seq and microRNA array expression data.

To calculate the ASE score for *ACSL1*, heterozygous SNPs in the coding region of *ACSL1* gene were identified for each individual (3–6 SNPs per sample). The ASE score of an mRNA-Seq sample was calculated as the average value of absolute differences between the fraction of the reference allele and the fraction of the alternative allele for all heterozygous SNPs with at least a 10× coverage.

### WGBS data analysis

Paired-end reads from WGBS were aligned to the GRCh37-lite reference genome by using BSMAP (version 2.74), with the following parameters (-z 33 -f 5 -g 3 -r 0 -m 17 -x 600 -u). The average genome-wide bisulfite conversion rate was measured as the genome-wide CpH conversion rate. Raw methylation values including *β*-values 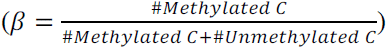 and *m*-values 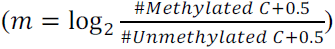 at individual CpG sites and SNPs were called by Bis-SNP (version 0.82.2)(Liu et al. 2012). Only CpGs with a coverage ≥5× were included for further analysis(Ziller et al. 2015). Because of the expected coverage differences between males and females as well as the DNA methylation changes during the X-inactivation process in females, we only analyzed autosome CpGs in this study.

Differential single CpG analysis was performed for CpGs surveyed in all samples, using a linear model with stepwise backward selection:

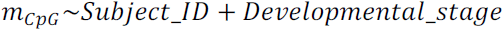

CpGs not showing strong inter-individual variations (i.e., difference between average *β* values of any 2 individuals <0.1) were excluded from inter-individual analysis. CpGs showing no strong developmental variations (i.e., difference between average *β* values of any 2 stages <0.1) were excluded from developmental analysis.

The sample-level methylation block pattern was established by regression tree–based segmentation(Chen et al. 2015) on *m*-values. Briefly, we assumed that *m*-values of CpGs in a segment are derived from a normal distribution that with a mean at its average *m*-value. The segment boundaries were defined by recursive partitioning–based regression-tree algorithms. We selected the optimal model that minimizes the Bayesian information criterion(Schwarz 1978):

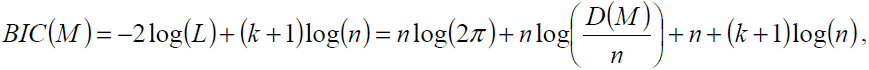

where *M* is a model of *k* segments from *n* observed CpGs. *L* and *D(M)* are the likelihood and deviance of the model, respectively. Neighboring segments with an average *β* difference ≤0.1 or *P* > 1 × 10^−6^ were recursively merged. Unique breakpoints between adjacent segments were collected from all samples, and the collection of regions included in the DMR analysis was defined across samples. The preprocessing and segmentation codes are available at http://ftp.stjude.org/pub/software/methylation-analysis-code.tar.

The i-DMRs and d-DMRs (range of *β* values ≥0.1) were identified in a linear regression model between regional average *m*-values and both subject IDs and developmental stages, followed by a stepwise backward model selection:

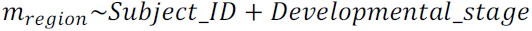

Regions with significant inter-individual or developmental differences (*q*-value ≤ 0.05) were classified as i-DMRS or d-DMRs, respectively.

Hierarchical clustering was performed by using the Spearman rank correlation distance on *β*-values from the 50,000 most variable CpGs or regions.

Correlation between blocks with variable methylation (range of *β* values ≥0.3) and adjacent genes with variable expression (methylation region overlapping 10-kb upstream of the TSS to the TES (at least a 2-fold difference between highest and lowest FPKM, with highest FPKM being ≥1) were calculated. To reduce potential over-segmentation from pooling breakpoints from all samples, gene or block pairs with *q*-value ≤ 0.5 were retained for merging, where adjacent blocks (no more than 500-bp apart) for the same gene with the same orientation (positive or negative) of correlation were merged. Correlations were re-evaluated, and pairs with final *q*-value ≤ 0.1 were considered differentially methylated regions significantly correlated with gene expression.

The *cisR*-mQTLs were detected by evaluating the association between *m*-values of the CpGs in the block and the genotype of SNPs within the region or up to 1000 bp away by using a linear model followed by stepwise model selection:

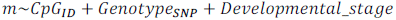

Adjacent blocks were merged by using the approaches described earlier. Regions with CpGs significantly associated with SNP genotype (*q*-value ≤ 0.05) were classified as *cisR*-mQTLs.

## Acknowledgements

We thank Dr. Vani J. Shanker for editing the manuscript. This study was supported by St. Jude Children’s Research Hospital–Washington University Pediatric Cancer Genome Project, Cancer Center support grant P30 CA021765 from the US National Cancer Institute and the American Lebanese Syrian Associated Charities of St. Jude Children’s Research Hospital.

